# The cost of multiplexing: PFC integrates multiple sources of information in non-orthogonal components accounting for behavioral variability

**DOI:** 10.1101/2022.12.28.522139

**Authors:** Julian L Amengual, Fabio Di Bello, Sameh Ben Hadj Hassen, Corentin Gaillard, Elaine Astrand, Suliann Ben Hamed

## Abstract

The frontal eye field (FEF) is a cortical area classically associated with spatial attention, perception, and oculomotor functions. FEF exhibits complex response properties through mixed selectivity neurons, allowing a high dimensional representation of the information. However, recent studies have shown that FEF encodes information in a low-dimensional regime hence limiting the coding capacity of the neural population. How the FEF encodes multiple sources of information with such limited encoding capacity remains elusive. To address this question, we trained two macaques to perform a visual attention task while we recorded FEF neuronal activity using multi-contact electrodes. FEF neurons encoded task- (time in the trial; CTOA) and behaviour- (reaction time, RT; focus of attention, TA) related parameters prior to the target onset. We found a clear modulation of the RT and TA as a function of the CTOA. Using dPCA, we characterized the functional relationship between neural populations associated with each parameter and investigated how this functional relationship predicts behaviour. We found that CTOA variability was associated with two different components the activation of which was correlated with the TA and the RT, respectively. These CTOA-related components were non-orthogonal with the RT and TA-related components, respectively. These results suggest that, when different sources of information are implemented during task performance, they show a very precise geometrical configuration in non-orthogonal components, which allows a high capacity of information coding at a cost of modulating both the capacity of the monkey to use attention information and its responsiveness toward external stimuli.

## Introduction

One fundamental question in neuroscience is how the brain organizes multiple sources of information to produce behaviour. In this context, it is essential to identify which components explain variability in performance, how these components interact at the neural level and how cognition emerges from this interaction. Given its rich connectivity with cortical sensory and high-order association areas, and subcortical structures associated with different cognitive aspects such as memory, the prefrontal cortex (PFC) is a brain region that crucially codes and combines multiples sources of information such as timing, motor feedback or the physical attributes of stimuli during task performance (1–5). Neurophysiology studies have revealed that the PFC holds neurons that exhibit complex response properties that reflect *simultaneously* the coding of different task- and behaviour-related parameters, a property that is called mixed-selectivity(6–9). This property plays a crucial computational role because it is related to the dimensionality of the neural representations (7), as these cells allow a high dimensional representation of the information in orthogonal (i.e. mathematically independent) components. This is considered as an advantage both in facilitating the readout of the information of interest and in implementing a larger number of cognitive computations. On the other hand, it has been proposed that neural population activity in the prefrontal cortex is low-dimensional, as the brain activity only visits a small fraction of all its potential states, hence limiting the coding capacity of the neural population(10–12). An open question in the field is thus how the PFC encodes multiple sources of information with such a limited capacity of encoding imposed by the low-dimensional representation of the activity. To address this, it is necessary to have access to the low-dimensional representation of the neuronal populations and to study whether these components are associated to the specific sources of neuronal variance and how these components interact to produce specific patterns of behaviour. In other words, the question is to what extent the coding of different parameters is performed by orthogonal representations of the information in the PFC and, if not, what are the behavioural implications of the non-orthogonalization.

The low-dimensional structure of the neural recordings can be analyzed using dimensionality reduction methods (13). In particular, demixed PCA(14) allow the extraction of components the variance of which is linked to specific parameters related to the task or to the subjects’ behaviour without imposing the orthogonalization constraint(11,15) (in contrast to the PCA where principal axes are all orthogonal). In a recent study, we used dPCA to show that the decoded position of attention in space and the behavioural outcome lie in different but non-orthogonal components. The degree of non-orthogonality varied from one session to the next and accounted for the behavioural gain associated with the prefrontal implementation of attention: the more orthogonal these components were, the higher the behavioural gain associated with prefrontal attentional function. This observation suggests that non-orthogonality might be a strategy of the PFC to encode multiple sources of information and might come at the cost of with a functional and behavioral interaction between these sources.

In the present study, we analyzed electrophysiology data from intracranial recordings in the frontal eye field (FEF, bilaterally) of two monkeys performing 100% validity-cued visual attention task. We show that the recorded cells encode multiple sources of information simultaneously accounting for the instructed attention position, the attentional focus (i.e. the spatial precision with which attention instruction was implemented), the time in the trial and the reaction time. Using dPCA, we identify the demixed components accounting for the variance associated with these sources of information. We show that the components associated with the time in the trial are non-orthogonal to the components associated with both the reaction time and the attentional focus. In addition, we show that each of these components accounts for behavioural variability and interaction between components. Overall, this work provides evidence of the rich capacity of the FEF to encode multiple sources of information in non-orthogonal representations, and describes the behavioural implications of such coding strategies.

## Results

Two rhesus macaque monkeys (M1 and M2) performed a 100% validity spatially cued target detection task consisting in responding to a change in luminosity of a spatially pre-cued landmark located in one of the four quadrants of a screen (Figure 1A), receiving liquid rewards for correct detection. In ∼60% (iqr 3) of the trials, monkeys were exposed to one or several changes of luminosity of uncued landmarks during the cue-to-target time interval (CTOA). These acted as distractors and had to be ignored. When monkeys responded to these distractors, the current trial was considered a False Alarm (FA) and they were not rewarded.

**Figure 1.**
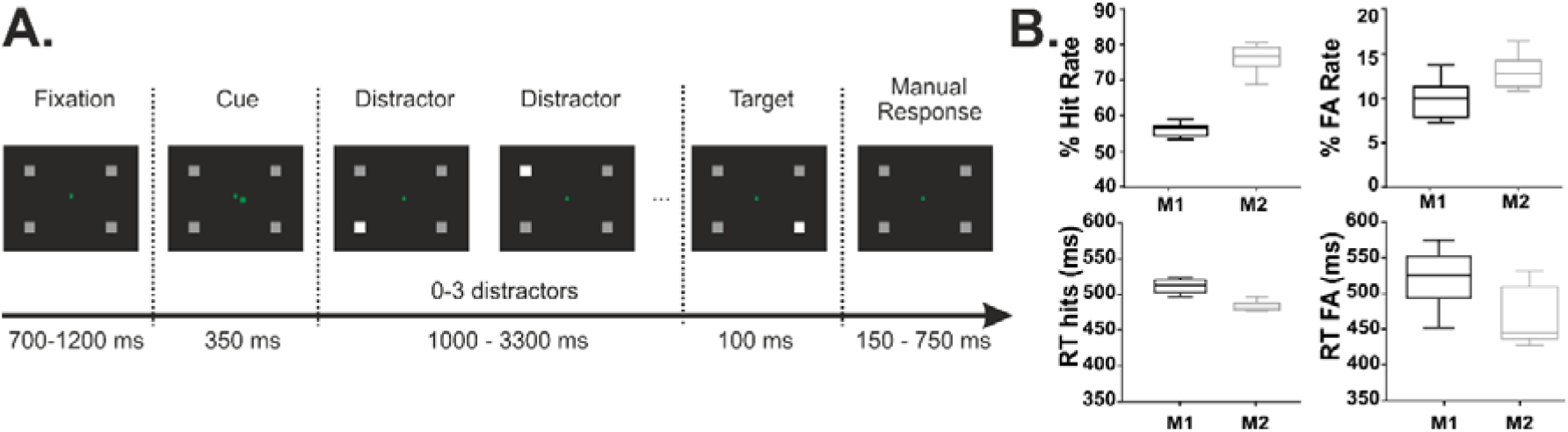
(A) 100% validity cued target-detection task with distractors. In order to initiate the trial, monkeys had to hold a bar with the hand and fixate their gaze on a central cross on the screen. Monkeys received liquid reward for releasing the bar manually 150-750 ms after target presentation onset. Target location was indicated by a spatial cue presented centrally (green square). Monkeys had to ignore any uncued event (distractors). (B) Barplot showing the percentage of hit rate, the percentage of false alarms, reaction time of hits and reaction time of false alarms of each monkey (M1 and M2) separately.

Monkeys M1 and M2 achieved 55.6% and 76.2% correct responses, respectively. Median reaction times of these responses were 502 ms (IQR: 14) for M1 and 472 ms (IQR: 7) for M2. FA rate to distractors was 10.5% (iqr 3.7) for M1 and 13.4% (iqr 6) for M2. Median reaction times of FA responses were 524 ms (IQR: 63) for M1 and 439 ms (IQR: 78) for M2 (Figure 1B). This type of behaviour is classically taken as an indication that the cue provides monkeys with advanced knowledge about relevant task items, allowing them to orient their spatial attention towards the expected target location(16–18), enhance their perceptual sensitivity(19,20) and prepare their behavioural response(21,22). In the following, we further show that this task imposes the interaction between different cognitive processes (attention, temporal expectation and motor readiness) all of which are encoded in the FEF and contribute to overall neuronal variance.

### Selective spatial attention is encoded in the FEF

MUA was obtained from 848 contacts (over 18 recording sessions, therefore 48 contacts implanted per session) implanted in the FEF (Figure 2A), a structure that is known to play a key role in covert spatial attention(23–25), while monkeys were performing the cued detection task. Overall, we found that 68.4% of all units had a significant activation to both the cue and target onsets (Wilcoxon paired test, p<0.05), and 65% of these active units showed a higher spiking rate in the post-target epoch ([0-400] ms) than in the pre-target epoch. Reproducing previous reports on FEF neuronal responses(22,26–28), we studied task-related modulations of MUA responses by comparing neuronal activity when attention was cued to the most preferred or the least preferred visual quadrant of each neuron. Figure 2B shows that the spiking rates were higher when the preferred location was cued (time period 100 ms to 400 ms post cue onset, Figure 2B, left). This higher firing rate regime associated to the preferred cueing location was sustained along the even during the target onset (Figure 2B, right).

**Figure 2.**
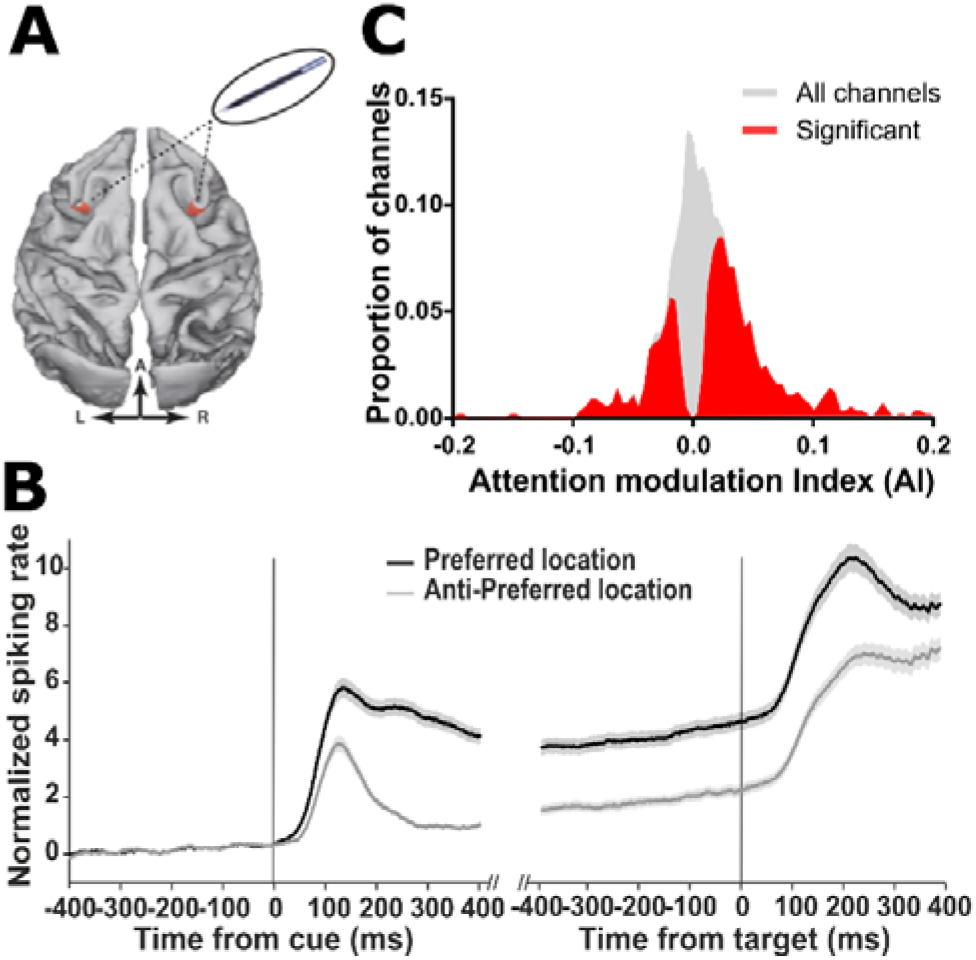
Location of electrophysiological recordings and characterization of their attentional modulation. (A) On each session, one 24-contact recording probe was placed in each FEF. (B) Single MUA mean (± s.e.) locked to the cue onset (left) and target onset (right) associated to when cue is orienting attention towards the preferred (black) or the anti-preferred (gray) spatial locations (C) Distribution of attention modulation index (Preferred - Anti-preferred)/(Preferred + Anti-Preferred), computed over 200 ms before target onset across all MUAs of all sessions. Red histogram corresponds to channels in which the neuronal activity during this time interval was significantly different between the preferred and the anti-preferred spatial attention responses (Wilcoxon test, p<0.05).

Firing rate differences between the preferred and the anti-preferred quadrant were converted into a modulation index, ranging from -1 to 1. Positive (negative) values of this index indicated higher activity at the preferred (anti-preferred) location. Median modulation index of the entire population was 0.008 (65% of units showing significant modulation, median significantly modulated units: 0.023, Figure 2C in red) and the distribution of index values was significantly greater than zero (p<.0001, Wilcoxon paired test) (Figure 2C). In addition, we tested whether the FEF encoded the cue position (selective spatial attention). Supplementary figure 1A shows the firing rates averaged across all MUA channels (n=848) locked to the cue, per each cue position. At the postcue interval of [0-400 ms], 75% of cells showed cue-position tuning. Overall, these results confirm that the recorded activity is sensitive to spatial attention, a well-documented property of FEF neurons.

### FEF encodes time-expectation, focus of attention and motor response-related information

In addition to the cued spatial attention, we investigated whether other sources of information related to the task (time expectation) and behaviour (the focus of attention in space and response times) were encoded in the FEF. To address the time expectation, we used the randomized time interval between the cue and the target onset (the cue-to-target onset asynchrony, CTOA) as a measure of target onset expectation. The distribution of the CTOA is shown in figure 3, upper panels. We observed that the distribution of the CTOA is not unimodal (Kolmogorov-Smirnov test, p <0.001). We measured the number of modes of distribution of the CTOA using a smooth bootstrapping method (number of repetitions = 5000) described in (29), which consists in testing parsimoniously whether the distribution of CTOA shows an increasing number of modes until reaching significance. We found that the distribution of the CTOA showed five significant modes (p = 0.04) at CTOA = 1080 ms, 1400 ms, 2000 ms, 2600 ms and 3000 ms. Reaction times (RT) were calculated as the time passed between the target onset and the manual response onset. In order to get rid of early responses, we applied the linear approach to threshold with ergodic rate (LATER) model (30) and we eliminated those reaction times that were below the early response upper threshold. The distribution of the resultant RTs is shown in figure 3 (intermediate panels, Median = 464 ms, IQ = 68). The focus of attention was calculated as the distance between the decoded position of the attentional spotlight to the real position of the target (see methods and (18,26,31–33)). Therefore, the focus of attention is defined as the Target-to-Attention distance (TA) measured shortly before the target onset. It has been previously described that this overt measure is linked with the probability to hit the target (18,22,31,34) and with the reaction time (31), accounting for how well subjects are orienting attention in space(22,31,32). The distribution of the TA is shown in figure 3 (lower panels, Median = 11.9°, IQR = 11.02).

**Figure 3.**
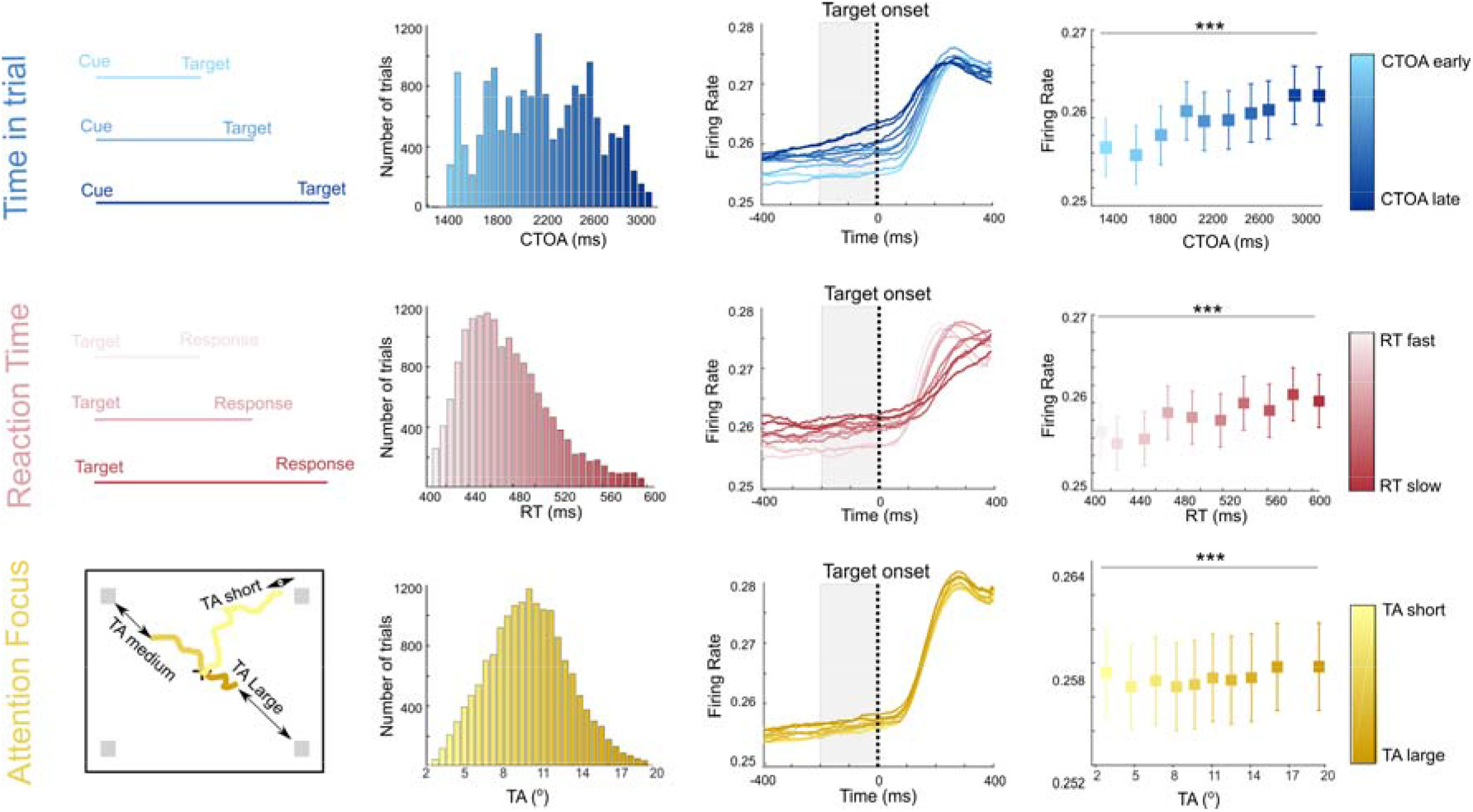
MUA activity in the FEF encodes time in trial, reaction time and attentional focus. Averaged MUA activity across all contacts (N = 848) grouped by time in trial (CTOA, blue), reaction time (RT, red) and attention focus (TA, yellow). CTOA reflect the time between the cue and the target, the reaction time reflect the time passed between target onset and manual response onset, and the TA is the distance between the decoded attentional spotlight and the target position 100 ms before the target onset. Histograms of these parameters across all trials are shown (middle left column). Tone colors represent the gradient of the parameter values (darker tones, higher values) for each of the parameters. The evolution of the time series MUA locked to the target onset for each parameter bin are shown (middle right column). Boxplots showing the median and the interquartile range of the MUA activity before the target onset (shaded rectangle in middle right columns) is shown (right column, Friendman test, *** p<0.001).

We studied the firing rate modulation as a function of these three different parameters. For each recording session, we sorted ascendingly the CTOA, TA and RT respectively and we binned these vectors in ten different bins (Figure 3, coloured from lighter shades to darker shades, middle right columns). We selected the trials based on each of these bins for each of these three parameters, focusing our analysis on hit trials. After equalizing the number of trials per each of the bins in each session, we obtained an average of 110 trials per bin (max = 149, min = 86). We averaged the firing rate (locked to the target onset) obtained from trials in each bin and parameter (Figure 3, right columns). Figure 3 shows that the firing rate was modulated as a function of the time expectation (CTOA; blue, upper panel), reaction time (RT; red, intermediate panel) and focus of attention (TA, yellow, lower panel). We found that FEF activity encoded these three different parameters (Friedman test, p<.0001 for all parameters). More specifically, we found that the firing rate increases linearly as a function of the CTOA (Spearman correlation, rho = 0.95, p < 0.0001) in the pre-target interval -200 to 0 ms. Similarly, in this same interval, the firing rate increased linearly as a function of the RT (Spearman correlation, rho = 0.92, p<0.001). However, firing rates showed a quadratic relationship with the TA (quadratic fit, R^2^ = 0.66, p =0.02) but not linear (Linear fit, R^2 = 0.23, p=0.15) in this same pre-target interval. The same results are reproduced for each monkey separately (Supplementary figure 2).

FEF shows mixed-selectivity for time-expectation, focus of attention and motor response related information. The neural mechanisms of prefrontal cortex neurons that allow encoding simultaneously multiple sources of information, a phenomenon called mixed selectivity, has been well studied in the literature (6,35,36).

Accordingly, we investigated whether the activity recorded in the FEF for these three different parameters, time in the trial, reaction time and attentional focus, showed mixed selectivity between these parameters and how this mixed selectivity fluctuated in time. Figure 4A shows that the selectivity of the neural population for each parameter is dynamic. Overall, there is a higher proportion of position-cells compared with the proportion of cells tuned to the rest of parameters. However, this proportion changes between before and after target presentation. Specifically, there is an increase of proportion of position-cells after the target onset (73% before the target onset vs. 82% after the target onset), due to the fact that pre-target epoch identifies spatial attention information while post-target epoch identifies visual responses to the target. The same effect is observed for RT-cells (37% before the target onset vs. 65% after the target onset). In contrast, the proportion of time-estimation cells tends to decay during the post-target interval (45% pre target vs. 32% post-target). The proportion of TA-cells remained constant irrespective of the target onset. We measured the percentage of neurons that showed random tuning. To do this, we randomly selected 10 subsets of trials, and we assigned them blindly to random CTOA, RT, position and TA conditions. We then measured the percentage of neurons that showed significant tuning (using non-parametric Friedman test). After repeating this process 1000 times, we selected the 95^th^ percentile of the distribution of the percentages of tuned cells. Using this approach, only 6% of cells showed significant random tuning, indicating that the tuning of the neural population to all these parameters was significant. We also measured the proportion of cells that were tuned *only* to one single parameter, just before the target onset (−100 ms to 0). We found that 27.2% of selective cells were tuned to only one parameter (17.7% to the cued position, 2.1% to the TA, 2.7% to the RT and 4.5% to the CTOA). In the following, we sought to investigate the proportion of cells that showed mixed selectivity in the pre-target interval that is during the period in which the attention information was held (Figure 4A, shaded area). Per each pair of parameters, we measured the proportion of recorded units that were tuned to each of the parameters (simple selectivity), to both parameters simultaneously (mixed selectivity) and to any of the two parameters (non-selective cells). The results are shown in figure 4B. We observed that per each pair of parameters we found a proportion of neurons that were tuned to both parameters simultaneously (minimum proportion, RT and TA, 6%; maximal proportion, Position and TA, 39%). We confirmed this rich and complex mixture of tuning selectivities by computing the proportion of cells that were tuned simultaneously to any combination of parameters (see Figure 4C). All in all, we found that the recorded neural population showed a complex pattern of mixed selectivities involving all these parameters simultaneously.

**Figure 4.**
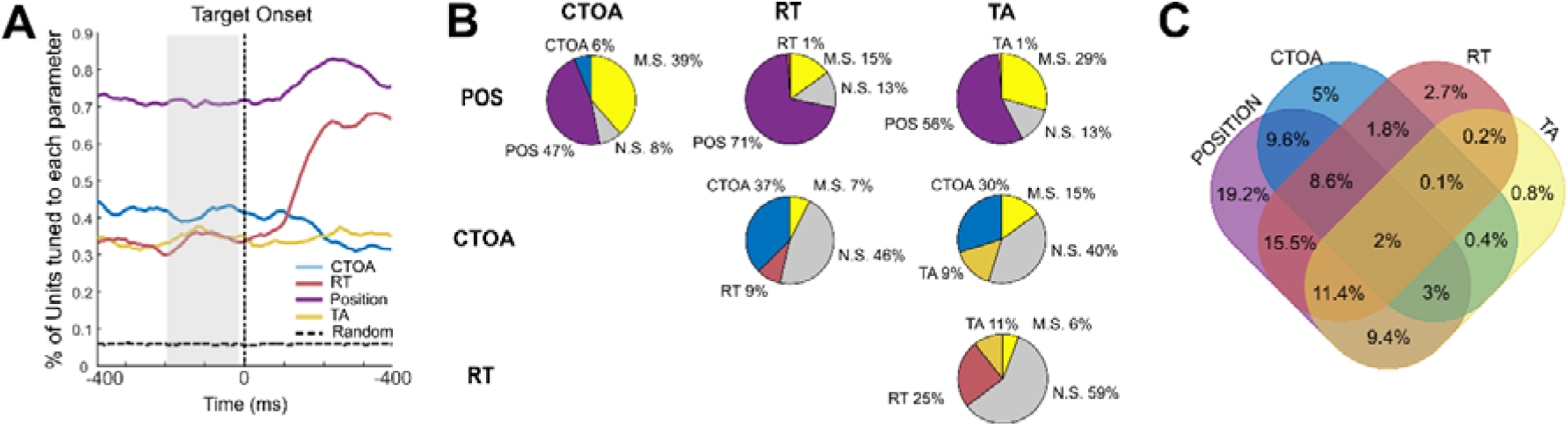
Mixed selectivity in the FEF. **(A)**. Time series reflecting the selectivity of the cells (measured as the proportion of cells) for each parameter (CTOA in blue, RT in red, TA in yellow and Position in purple) locked to target onset. Dashed line represents the 95^th^ C.I. of cells tuned to a random process. **(B)** Pie charts showing the proportion of cells showing no selectivity, single selectivity and mixed selectivity to each pair of parameters. **(C)** Venn diagram showing the proportion of selectivity for each combination of parameters. Percentages reflect the proportion of cells tuned to each of the intersections of tuning populations.

### Neural population holds a low-dimensional representation of Time-expectation, focus of attention and motor response in the FEF

Previous literature has reported evidence that the activity in the prefrontal cortex is contained in a low-dimensional manifold that can be extracted projecting the firing rates from the high-dimensional recording space onto few latent variables that represent the majority of the variance in the recordings(7,13,37). In the previous section, we have demonstrated the ability of the PFC to encode simultaneously different types of information. The question we address now is how the PFC organizes geometrically the representation of these different sources of information. To solve this issue, we have applied a demixed principal component analysis (dPCA)(14,38) which consist on a supervised dimensionality reduction method that decomposes (or demixes) the data in components by marginalizing it over different parameters. Differently to the principal components analysis where the variance is described in the direction of the eigenvectors of the covariance data matrix, with this model each component describe variance only in the direction of the eigenvectors of the covariance of each of marginalization.

First, we sought to unmix each of the parameters from the rest of parameter independent activity. Supplementary figure 1B shows how the dPCA succeeds in extracting specific position-related variance (65% of the total variance). We projected the firing rates corresponding to trials categorized as a function of each cue position presentation (activity locked to the target onset) onto the first three components that accounted together for 79% of the total variance in the data. The second and third component were associated to position-related variance, whereas the first component showed the task-related variance independent on position. Therefore, this analysis shows that the recorded MUA activity in the FEF might be confined in a three dimensional space containing position information (dimensions two and three) and task-related dynamics in independent axis (dimension 1).

In the following, we investigated whether we could reconstruct a low-dimensional representation of the recorded neural population the axis (or neural mode) of which represents variance specific for each parameter of interest (time-expectation, focus of attention and motor response). Demixed PC analysis showed that 23% of the variance in data was explained by time expectation (Figure 5A, top). Firing rates locked to the target onset (−400 ms to 400 ms) were projected onto the third and fourth demixed principal component (associated with CTOA). These two components showed a different pattern of dynamics as a function of the CTOA bin (Figure 5A, bottom). MUA activity projected onto the component 3 (8.2% of variance) showed that activity regime changed linearly (trial level projection, Linear Fit, R^2 = 0.98, p = 2e-14) as a function of the CTOA. Indeed, the higher the time between the cue and the target, the larger was the firing rate projected onto this component. A different pattern was observed when MUA activity was projected onto the forth demixed component (2.2% of variance). Activity projected onto this component showed a quadratic relationship between the normalized firing rate and the CTOA (trial-level projection, quadratic fit, R^2 = 0.97, p = 3e-15). Specifically, activity was maximal at CTOA bin = 2000 ms. We used a stratified Monte Carlo leave-group-out cross-validation (100 iterations) to test whether these components decoded the CTOA bin. We found that the accuracy of decoding CTOA was significantly above-chance level before target onset until 200 ms after the target onset (permutation test, 100 shuffles, p <0.05). These results were fully reproducible per monkey (supplementary figure 3). These results showed that time expectation was contained within a two-dimensional space, components of which represent the time-expectation through two different functional mechanisms.

**Figure 5.**
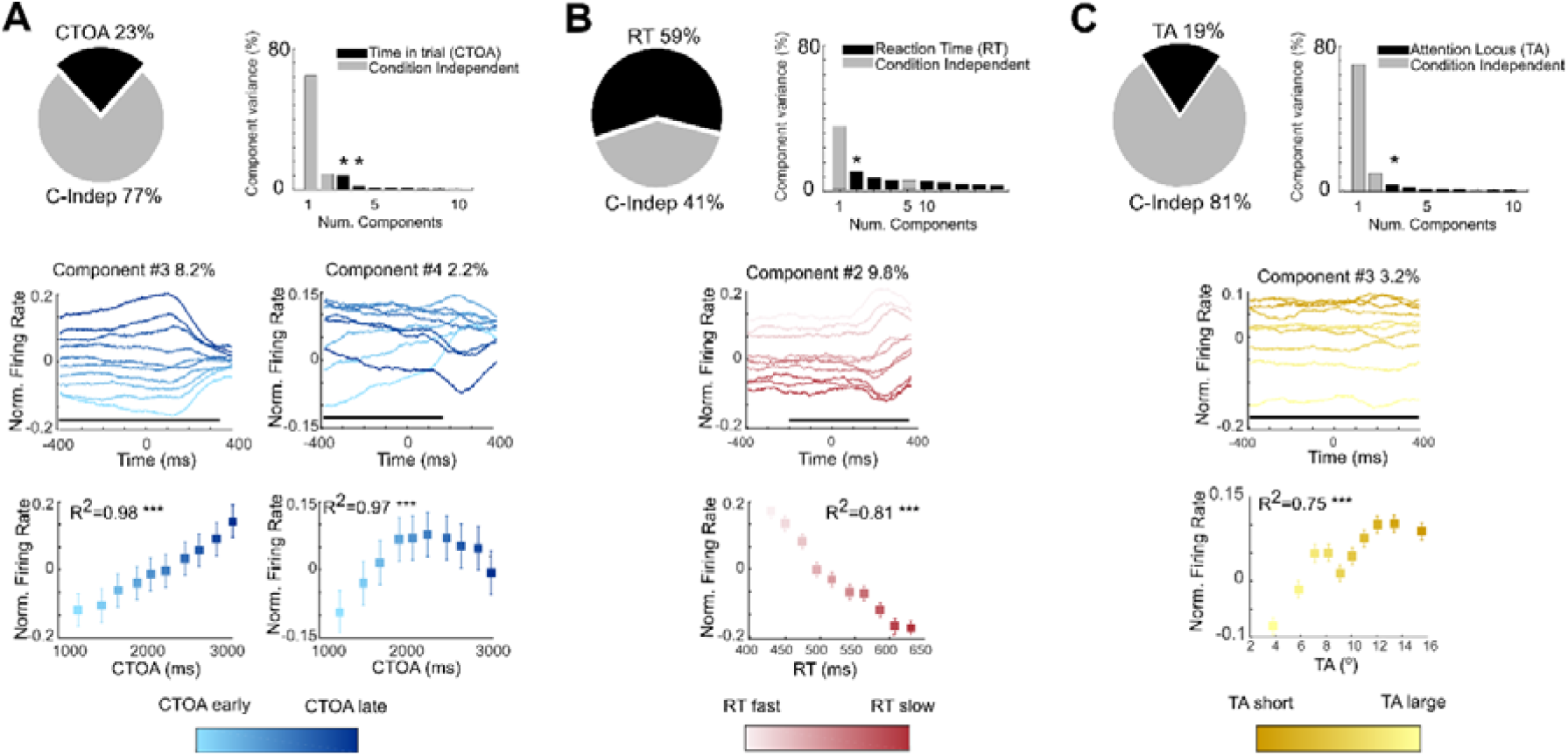
dPC analysis of each parameter. Demixed PC analysis reduces the dimensionality of the recordings and extracts variance specific of CTOA (A), RT (B) and TA (C). Pie chart represents the variance distributed between parameter-specific (black) and parameter-independent (gray). Bar plot shows the % variance (parameter specific, black; parameter independent, gray) attributed to the first 10 demixed components. Stars represent the parameter-specific components that decode the parameter at trial level above chance level. Only those are represented in time. Time series reflect the projection of the data onto the decoding parameter-specific components for each dPC analysis (CTOA, blue; RT, red: TA; yellow). Tone colors represent the gradient of the parameter values (darker tones, higher values) for each of the parameters. (*** p<0.001, regression model)

Similarly, we conducted the same analysis for the focus of attention and reaction time. For RT, dPCA showed that the 59% of the variance was linked to the reaction time (Figure 5B, top). MUA activity projected onto the component 2 (9.8% of variance) showed that activity regime changed linearly (trial level projection, Linear Fit, R^2 = 0.81, p < 0.001) as a function of the RT, the projected firing rate onto this component decreasing as a function of the RT (Figure 5B, bottom). We found that this component significantly decoded RT between 200 ms before the target until 400 ms after the target (permutation test, 100 shuffles, p <0.05). No other component showed decoding accuracy higher than the chance level. Therefore, dPCA found one single component that contained the variance linked to speed of response. Finally, dPCA showed that 19% of the variance was linked to the focus of attention (Figure 5C, top). MUA activity projected onto the component 3 (3.2% of the variance) showed that the projected firing rate increased as a function of the TA (Figure 5C, bottom, trial level projection, Linear Fit, R^2 = 0.75, p < 0.01). We found that this component significantly decoded TA bin during -400 ms to 400 ms with respect to the target onset (permutation test, 100 shuffles, p <0.05). No other component showed decoding accuracy higher than the chance level. Therefore, dPCA found one single component that contained the variance linked to the focus of attention.

### CTOA, RT and TA interact both in the neural low-dimensional representation and in behavior

In the previous section, we described low-dimensional representations in the neural population associated to parameter specific variance in the data. The next question is to what extend these representations are independent one from each other or to what extend they overlap. If they overlap, the expectation is that the angle between the neural modes corresponding to each of these parameters are found to be non-orthogonality. In order to measure this angle, we conducted a dPC analysis with the aim to obtain simultaneously different neural modes associated to CTOA, TA, RT and trial progression (independent from) the other three parameters. Demix PCA succeeded in reconstructing a low-dimensional representation of the data. More specifically, the 70% of the total variance of data was represented in the first 10 demixed principal components (Figure 6). Most of the variance was explained by trial-progression (46 %), followed by RT (22%), CTOA (17%) and TA (16%) (Figure 6B). Firing rates projected onto the first principal components of each parameter showed similar dynamics as described in the previous section (Figure 6A). Importantly, we found that these components were non-orthogonal, except for the RT and TA components (Figure 6C).

**Figure 6.**
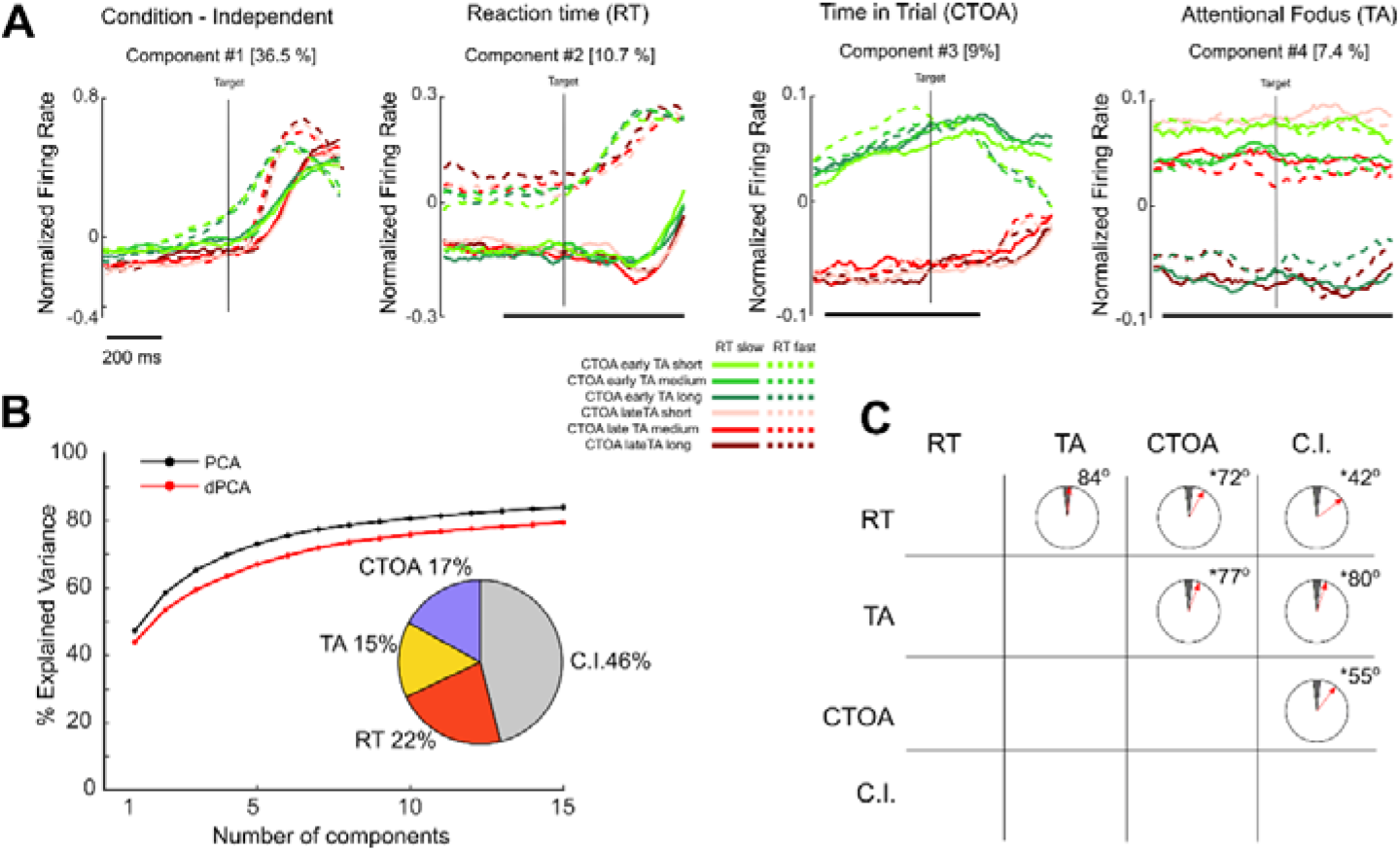
Low-dimensional representation of MUA recordings in a parameter-based space. **(A)** Time series corresponding to the projection of the MUA recording onto the condition-independent, reaction Time (RT), time in trial (CTOA) and attention focus (TA) specific components extracted simultaneously from a single dPCA. Red and green colors represent CTOA early and late (respectively), continuous and dashed lines represent fast and slow RT (respectively), and tones represent close (brighter), medium and long (darker) TA. Time series are locked to the target onset. **(B)** Cumulative signal variance explained by PCA (black) and dPCA (red) across the first 15 components. dPCA explains similar amount of variance than PCA. Pie chart shows how the total signal variance is split between CTOA, TA, RT and condition independent. **(C)** Representation of the angle between pairs of axes. Dark section reflects the angle values where angle values are considered statistically orthogonal. Red arrow indicates the actual angle between each pair of axes. Stars mark the pairs that are significantly and robustly non-orthogonal.

Thus, the low dimensional representations associated with CTOA and RT and TA, respectively are not orthogonal. The next question we wanted to answer was whether there was a behavioral interaction between these three parameters. To this aim, per each session, we divided the trials into 10 different bins based on their CTOA, and we measured the median RT and TA within each bin. We found a clear linear relationship between the CTOA and the averaged RT (averaged value per all sessions, linear regression R^2 = 0.84, p<0.001) (Figure 7A), whereas the relationship between the CTOA and TA was quadratic (Linear regression, R^2 = 0.07, p=0.4; quadratic regression, R^2 = 0.59, p<0.05, Figure 7B). In addition, we found that CTOA and TA also affected the likelihood to respond to the target onset (Figure 7C and 7D, Linear regression between CTOA and hit rate, R^2 = 0.82, p<0.01; Linear regression between TA and hit rate, R^2 = 0.82, p<0.01). Finally, we observed that the average reaction time and TA over each CTOA bin highly correlated with the level of activation of the first and second demixed principal components associated with the CTOA variability, respectively (Linear Regression between RT and dPC1, R^2 = 0.79, p<0.01; Linear Regression between TA and dPC2, R^2 = 0.49, p<0.01). As described in Astrand and colleages (31), we replicated that monkeys responded faster when their attention was close to the target onset (Supplementary Figure S4). These results strongly suggest that changes in the RT and TA across CTOA correspond with specific levels of activation of the neural population variability linked with the CTOA.

**Figure 7.**
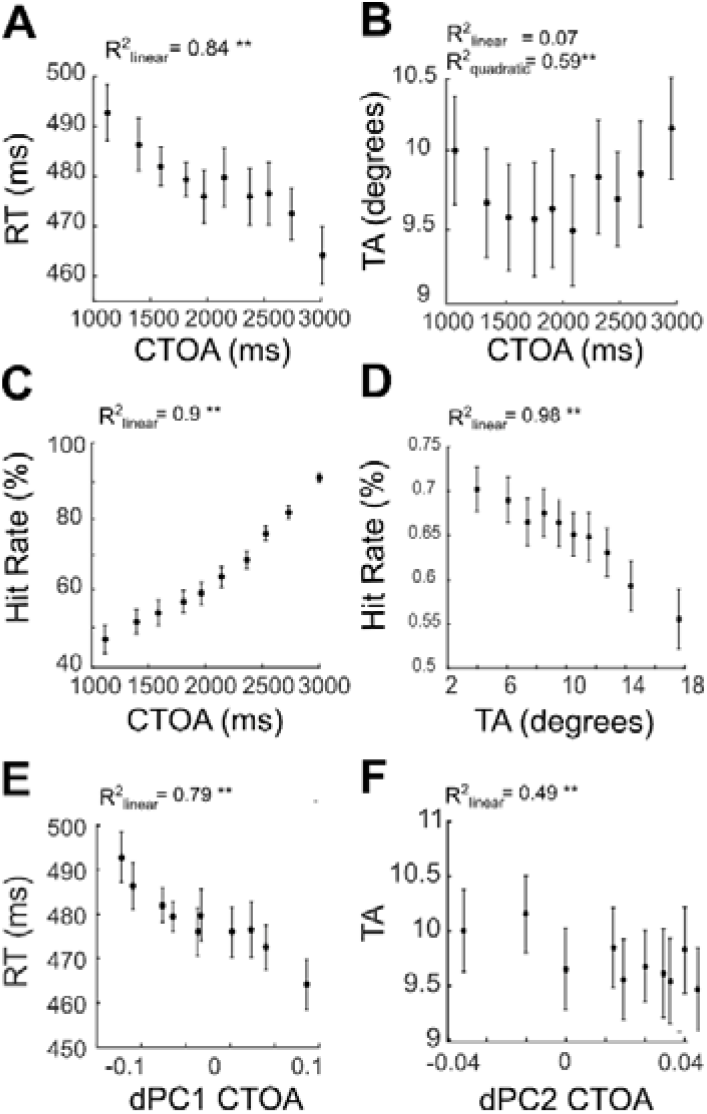
Behavioral results. **(A)** RT as a function of CTOA averaged across sessions. Each square represents, per each CTOA bin, the mean RT value. Bars represent the standard error of the RT measure across sessions. **(B)** TA as a function of CTOA averaged across sessions. Each square represents, per each CTOA bin, the mean TA value. Bars represent the standard error of the TA measure across sessions. **(C)** Hit rate as a function of CTOA averaged across sessions. Each square represents, per each CTOA bin, the mean hit rate value. Bars represent the standard error of the hit rate measure across sessions. **(D)** Hit rate as a function of TA averaged across sessions. Each square represents, per each TA bin, the mean hit rate value. Bars represent the standard error of the hit rate measure across sessions. (E) RT as a function of the activity projected onto the first dPC associated to CTOA across sessions. For each session, and each CTOA bin, the mean projected firing rate at the time interval -300 to 0 ms previous to the target onset as well as median RT for those trials. Average RT across sessions (square symbols) and s.e. were plotted against average firing rate. (F) Same as in (E) but for TA. (* p<0.05, **p<0.001 regression model).

## Discussion

In the present work, we have shown the FEF ability to encode simultaneously multiple sources of information in latent, non-orthogonal components and we have described how such computational organization explains specific behavioral interactions between these parameters. More precisely, we have analyzed electrophysiological data recorded bilaterally from both FEF in two macaque monkeys performing a 100% validity cued attention task. In each trial, they were presented with a cue located in one of the four quadrants of the screen, and they were instructed to respond to a target stimulus located in the same quadrant after a randomized time interval (CTOA) while avoiding distractors. Using a linear regression algorithm (18,31,33,39), we have decoded the position of the attentional locus in the space and estimated the distance between this decoded position of attention to the real target position before target onset (Attentional focus, TA). We divided our trials based on four different parameters that accounted for the CTOA, TA, response time (RT) and the cued position. We found that the firing rate regime on the FEF, measured before the target onset, differed based on each of these parameters, indicating that the FEF encoded these four different sources of information. In addition, we classified the recorded cells based on their selectivity to these parameters. While we found cells showing a very high single specificity to one parameter, we found cells showing mixed selectivity to more than one of these parameters. Using demixed PCA(14), we found that the variance in our recordings could be split in different components the variability of which was explained by these parameters. The components associated with CTOA were non-orthogonal with the components linked with the variability associated to RT and TA, respectively. Finally, behavioral analysis revealed that RT and TA changed as a function of the CTOA, and in correlation with the amount of activity of the neural states associated with the CTOA. Overall, these results show the high coding capacity of the FEF that can be attributed to concurrent multiple neural processes encoded using non-orthogonal representations that account for overt behavioral interactions.

At neuronal level, we have shown that the average firing rate of the recorded neural population increases as a function of the CTOA. This is in line with previous studies showing an increase of the firing rate as a consequence of a reward expectation(40–42), or the effect of an iteration of recurrent processes along the cue to target interval, which contains the representation of when (temporal expectation) and where (spatial attention) the target will appear(43). All in all, our results replicate previous findings suggesting an upper-modulation of the activity in the FEF during the cue to target interval. We also found a clear modulation of the firing rate before target onset as a function of the reaction time, such that faster responses were preceded by a higher firing rate than slower responses. Previous literature has shown that reaction times were reliably associated with the amount of time required by FEF cells to reach a certain firing rate threshold in saccadic(28,44–46) and manual responses(47). Concretely, these studies show that FEF neurons must be depolarized more vigorously for initiating responses with stronger input in order to speed up response times. This indicates that reaction times are sensitive to the modulation of the activity in the FEF, in agreement with our observation. Finally, we found that FEF cells were tuned to the attentional focus, understood as the distance between the decoded position of the attentional spotlight to the target onset, measured before the target onset(22,27,31,34). This is in line with recently reported evidence showing that attention is dynamic and rhythmic in space(27,48), and that when pooling trials based on attention to target distance, spiking rate of FEF cells increased as closer was the attentional spotlight to the expected position of the target(17,22).

Importantly, we found that the majority of the recorded neurons on the FEF encoded these parameters through mixed selectivity cells. More specifically, a 72% of the recorded cells showed tuning to more than a single parameter. These mixed selectivity cells have been reported in different brain areas(6,36,49,50), including the FEF(22,35), and they represent a signature of high-dimensional neural representation of relevant information and task-related parameters (7). Particularly, our results show that certain recorded cells are tuned simultaneously to more than two parameters, namely, spatial attention, focus of attention, RTs, and CTOA. To our knowledge, only one study has reported evidence of such capacity of mixing information in a neural population. Ledergerber and colleagues show simultaneous tuning of position, head direction and speed in cells recorded in the CA1 and subiculum in rats(51). They convey that such rich mixing of features in a specific neural population might indicate a high capacity of information transmission towards downstream regions, integrating simultaneously different covariates and ensuring that such wide number of covariates are accessible to distant projection areas. Similarly, the FEF has been considered in the literature a “*hub*” region, containing visual, motor and visuo-motor cells, projecting towards dorsolateral prefrontal cortex, cingulate, parietal and posterior cortices(44,45,52). Such rich anatomical connectivity might explain this multiple variety of tuning capacities for different task-parameters and behavior-related parameters as reported here.

We studied how much variance in our data was related to each of our parameters (CTOA, TA and RT) by using different dPCA specific to each of these parameters. For both RT and TA, we found that the dPCA successfully encoded each of the parameters in two components the activation of which showed a linear relationship with these parameters. For the CTOA, we found a more complex picture that deserve further discussion. Specifically, we found that the dPCA successfully encoded the time in trial in two different and orthogonal components, showing two different activation regimes. The projection of the activity associated with each CTOA bin onto the first demixed CTOA-component revealed a linear increase of the activity state, being low in the early stages of the trial and linearly increasing for longer CTOAs. We found that these states of activation nicely correlated with the reaction time measured in these CTOA-based selected trials, suggesting a functional link between this CTOA-related component and reaction time variability. This result accounts for the observed correlation between CTOA and RT, whereby responses performed in early stages in the trial were slower than those responses performed in the late stages of the trial. In contrast, projected firing rates onto the second demixed CTOA-component showed an inverted U-shape, in which lower activity state corresponded in early and late trials, while intermediate CTOA trials showed the maximal activation. When the activation of this component was the lowest (coinciding with trials selected at early and late stages of the trial), the encoded position of the attentional spotlight was more distant to the expected position of the target, while this distance was shorter when the activation of this component was the highest. This finding suggest a functional link between the CTOA-related variability explained by this component and the attentional focus. Prior studies have addressed the functional relevance of high-order, non-linear components, as the one observed by this CTOA-component. Recently, Okazawa and colleagues(53) suggested that these curved components might arise automatically due to fundamental constraints on neural computations that limit the dynamics of the firing rates, including the fact that expected values of the firing rate are non-negative, that they are bounded by metabolic constraints(54) (Lennie 2003), and that there is a strong evolutionary pressure to precisely encode information(55). Following the arguments of Okazawa et al. (53), we speculate that these CTOA-related components might define a sub-region in the state space where sensory information is encoded, defining a curved manifold subserving efficient readout from other brain regions (here, readout corresponding to efficient attentional orientation). This needs to be tested experimentally.

Finally, we applied a dPCA to simultaneously decompose our variance dataset in components associated to the CTOA, RT and TA. As reported in Kobak et al(14), this dimensionality reduction method does not assume the orthogonality of the components when efficiently linking specific sources of variance to each of the principal components. We studied the geometrical properties of the low-dimensional manifold containing the first principal components associated to each parameter in terms of the orthogonality between these components. We found that the CTOA component was significantly non-orthogonal to both the TA- and RT-related components. This result indicates that the FEF encodes these sources of information through non-orthogonal, partially overlapping codes. Previous literature has reported that encoding multiple sources of information in orthogonal codes through mixed-selectivity cells might ease the accessibility of the information to the downstream neurons(7,56). However, the low-dimensional manifold in which the recorded data is embedded limits the number of different encoding information that may be implemented in the system. It has indeed been reported that primate prefrontal neuronal populations have an informational capacity often limited to 3 sources of information(57). We argue that the FEF might therefore code multiple sources of information in non-orthogonal components as a strategy to maximize its encoding capacities. However, this strategy of encoding might come with a cost. We observe in our data a clear interaction between the CTOA and RT (responses get faster as CTOA advances) and CTOA and TA (attention is closer to the target in middle stages of the trial). Therefore, these results suggest that encoding through non-orthogonal components might drive the way the monkey uses attention and responsiveness as a dynamic cognitive resource to accurately perform the task. In support of this, we have recently reported evidence that the degree of non-orthogonality between the attentional component and a component which variance is associated with the reported behavior of the monkey, accounts for the degree to which the monkeys can use attentional information to guide behavior(22). In conclusion, we report converging evidence showing that different sources of information are implemented simultaneously in the FEF in a very precise geometrical configuration, in non-orthogonal components. This configuration is proposed to allow for high capacity of information coding, at the cost of modulating both the capacity of the monkey to use attention information and its responsiveness toward the stimulus. This finding sheds light onto the dynamic changes of the computational mechanisms of the attentional system during task performance, and it is expected to have profound implications in the development of efficient decoding algorithms aimed to extract specific cognitive information from electrophysiological recordings.

## Material and Methods

### Endogenous cued detection task and Experimental setup

The task is a 100% validity endogenous cued luminance change detection task (Figure 1A). The animals were placed in front of a PC monitor (1920×1200 pixels, refresh rate of 60Hz) with their heads fixed. Stimulus presentation and behavioral responses were controlled using Presentation® (Neurobehavioral Systems, Inc.). To start a trial, the monkeys had to hold a bar placed in front of their chair, thus interrupting an infrared beam. The appearance of a central fixation cross (size 0.7°×0.7°) at the center of the screen, instructed the monkeys to maintain their eye position (Eye tracker - ISCAN, Inc.) inside a 2°×2° window, throughout the duration of the trial, so as to avoid aborts. Four gray landmarks (LMs size 0.5°×0.5°) were displayed, simultaneously with the fixation cross, at the four corners of a hypothetical square having a diagonal length of ∼28° and a center coinciding with the fixation cross. The four LMs (up-right, up-left, down-left, down-right) were thus placed at the same distance from the center of the screen having an eccentricity of ∼14°. After a variable delay from fixation onset, ranging between 700 to 1200 ms, a 350 ms spatial cue (small green square - size 0.2°×0.2°) was presented next to the fixation cross (at 0.3°), indicating the LM in which the rewarding target change in luminosity would take place. Thus, the cue presentation instructed the monkeys to orient their attention towards the target in order to monitor it for a change in luminosity. The change in target luminosity occurred unpredictably between 750 to 3300 ms from cue onset. In order to receive their reward (a drop of juice), the monkeys were required to release the bar between 150 and 750 ms after target onset (hit). To test the monkeys’ ability at distractor filtering, on half of the trials, one of the two distractor typologies was randomly presented during the cue-to-target delay. In ∼17% of the trials (D trials), a change in luminosity, identical to the awaited target luminosity change, took place at one of the three uncued LMs. In these trials, the distractor D was thus identical in all respects to the expected target, except for being displayed in an uncued position. In ∼33% trials (d trials), a local change in luminosity (square) was displayed at a random position in the workspace. The size of the local change in luminosity was adjusted so as to account for the cortical magnification factor, growing from the center to the periphery (Schwartz 1994). In other words, d distractors had the same size as D distractors when presented at the same eccentricity as D. The absolute luminosity change with respect to the background was the same for both d and D. The monkeys had to ignore both distractor typologies (correct rejections – RJ). Responding to such distractors within 150 to 750ms (false alarm - FA) or at any other irrelevant time in the task interrupted the trial. Failing to respond to the target (miss) similarly aborted the ongoing trial.

### Electrophysiological recordings and spike detection

Bilateral simultaneous recordings in the FEF in both hemispheres were carried out using two 24-contact Plexon U-probes (Figure 1B). The contacts had an interspacing distance of 250 μm. Neural data was acquired using a Plexon Omniplex® neuronal data acquisition system. The data was amplified 500 times and digitized at 40,000Hz. Neuronal activity was high-pass filtered at 300Hz and a threshold defining the multiunit activity (MUA) was applied independently for each recording contact and before the actual task-related recordings started.

### Decoding procedure & attentional focus (TA) estimation

#### Training procedure

In prior studies, we showed that the endogenous orienting of attention (Figure 1C) can be reliably decoded from the FEFs activity using a regularized optimal linear estimator (RegOLE) with the same accuracy as exogenous visual information(18,22,27,33 for review,39,58). Here, we used the same approach to train a RegOLE to associate the neural responses prior to target onset ([-220 + 30] ms from target onset), based on a leave-one-out training/testing procedure, with the attended location, i.e., with the expected target presentation LM, based on cue information. Neural responses consisted in a vector containing the MUA signals collected at each of the 48 recording contacts during this pre-defined pre-target onset epoch. Our general objective here was to have as precise as possible an estimate of the attention position before a specific visual event, averaging activities over large enough windows to have a reliable single-trial estimate of the neuronal response on this window, while at the same time a not-too-large time window to have a reliable estimate of where attention was placed by the subject at a specific time in the task(22,27,34,59,60).

The RegOLE defines the weight matrix W that minimizes the mean squared error of **C*=* W* (R+b)**, where C is the class (here, four possible spatial locations), **b** is the bias and **R** is the neural response. To avoid over-fitting, we used a Tikhonov regularization(39) which gives us the following minimization equation **‖ W* (R+b)−*C*‖** ^**2**^**+ λ * ‖W‖**^**2**^.

The scaling factor *λ* was chosen to allow for a good compromise between learning and generalization. Specifically, the decoder was constructed using two independent regularized linear regressions, one classifying the *x*-axis (two possible classes: -1 or 1) and one classifying the *y*-axis (two possible classes: -1 or 1).

#### Testing procedure

In order to identify the locus of attention at the moment of target or distractor presentation in the 20 next new trials following the initial training set, the weight matrix defined during training was applied to the average neuronal activity recorded in the 150 ms prior to target. The described training (over 200 previous trials) / testing (over 20 novel trials) procedure was repeated after every 20 correct responses, by re-training the decoder with the new database composed by the last 200 correct trials. This continuous updating of the weight matrix W is implemented in order to minimize the impact of possible uncontrolled for changes in the recorded signal during a given recording session onto the decoding procedure.

### Estimating the (x,y) spatial locus of the attentional spotlight (AS)

The readout of the RegOLE was not assigned to one of the four possible quadrants by applying a hardlim rule, as usually done for classification purposes. Rather, it was taken as reflecting the error of the decoder estimate to the target location, i.e., in behavioral terms, as the actual (*x,y*) spatial estimate of the locus of the attentional focus to the expected target location(18). We show here and elsewhere(18,22,27) that this (*x,y*) estimate of the attentional spotlight (AS) accounts for variations in behavioral responses. In order to analyze how the distance of the decoded attentional spotlight to the target affected both behavior and neuronal MUA responses, we computed, for each target presentation, the distance between the decoded AS and the target (TA) as follows: 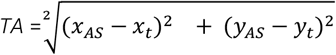 where ***x*_*AS*_** and ***y*_*AS*_** correspond to the Cartesian coordinates of the attentional spotlight (*AS*), and ***x*_*t*_** and ***y*_*t*_** correspond to the Cartesian coordinates of the target position (*T*).

### Partitioning of the neural data

Similarly to the decoding procedure, only neuronal data from correct trials were analysed. Neural data was partitioned based on four different parameters: 1) CTOA, i.e. time from cue onset and target onset; 2) RT, response reaction time, considered as the time between target onset and manual response and 3) TA or focus of attention, i.e., the distance between the decoded attention spotlight position and the cued LM position. For each of these three, we pooled trials in 10 different bins based on the deciles distribution of each parameter (Mean number of trials per class, 101 ±8). Trials in which the TA was longer than 18° were discarded.

### Behavioral Performance

Main task performance was measured through four different parameters: Hit rate (number of hits divided by the number of hits and misses), False alarm rate (Number of false alarms divided by the number of distractors), and the reaction time for hits and false alarms. Hit rate and reaction time to target responses were also calculated in each CTOA and TA bins.

We use a linear approach to threshold with ergodic rate (LATER) model to estimate the reciprobit distribution of latencies from responses to hits in the main task performance(30). This method was used to identify express responses from the distribution of responses in each session, and discard them for RT analysis.

### Characterizing MUA selectivity

In order to quantify the magnitude of the modulation of FEF individual to the orientation of the cue location, we pooled trials based on whether the cue oriented attention in the preferred spatial location within the neuron’s receptive field (RF), or non-preferred spatial location outside the RF, considering only correct trials. We defined a modulation index as:

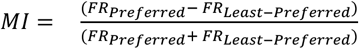 where ***FR*_*Preferred*_** and ***FR*_*Least−Preferred*_** corresponds to the median firing rate when the preferred and the least-preferred cue are presented. Firing rates were computed on the - 300 to -50 ms pre target epoch, z-scored with respect to a [-300 to -0] ms pre-cue epoch. The statistical significance of the MI was performed with a non-parametric Wilcoxon rank test (p < 0.05).

We measured the selectivity of each neuron for each of the parameters (CTOA, RT, TA and position) across the time interval -400 to 400 ms locked to the target onset. To this end, we calculated the main parameter effect across the interval (sliding window of 50 ms, step of 25 ms) by using a non-parametric Friedman test. With this method, we obtained per each time-subinterval the proportion of cells that were significantly tuned to each parameter. For each time point, we additionally measured the proportion of cells that showed tuning when trials were selected randomly. We repeated this process 1000 times obtaining a distribution of values corresponding to the proportion of cells that were tuned to random variability pear each randomization. For each time point, we selected the 95^th^ value of this distribution as threshold for random tuning.

Per each pair of parameters (CTOA-TA, CTOA-RT, CTOA-Position, TA-RT, TA-Position and RT-Position) we measured the selectivity of each cell for each of the two parameters (single selectivity) and for both parameters simultaneously (mixed-selectivity). To do this, we averaged the firing rate in the interval -300 ms to 0 ms before the target onset, and we used a non-parametric test Friedman test to measure the tuning for each parameter (p<0.05). A cell was considered to present single selectivity for a given parameter if its activity was tuned to this parameter and not tuned to the other parameter of the pair. A cell was considered to present mixed selectivity if it was significantly tuned to both parameters simultaneously. Cells without significant tuning to any of the two parameters of the pair were considered as non-selective.

### Demixed PCA

Recent research points that neural function is built on population activity patterns rather than on independent modulation of individual neurons (61). These patterns reflect the coordination of responses across neurons that corresponds to a specific neural mechanism underlying a specific behaviour (13). The population activity structure can be estimated by applying a dimensionality reduction technique to the recorded activity such as Principal Component analysis (PCA). Using this method, we can extract a number of latent variables (principal components) that capture independent sources of data variance providing a description of the statistical features of interest (13). However, this method does not take task- or behaviour-related parameters into account, mixing these sources of information within each of the extracted latent variables(14). With the aim of describing how much variance in the neural population can be explained by each parameter (CTOA, RT and TA), we performed a demixed principal component analysis (dPCA (14)), which captures the maximum amount of variance specifically explained by each of the above-defined parameters in each extracted latent variable and reconstructs the time course of the parameter-specific response modulation.

First, we described the amount of variance explained specifically for each parameter. To do so, we applied an independent dPCA analysis for each parameter, selecting trials based on the deciles of the distribution of the values in the parameter. In each of these analysis, the aim was to decompose the data into latent variables that estimate over time both the variance attributed to a parameter of interest and the variance independent to the parameter. Second, we aimed to describe how the different subpopulations associated to each parameter overlapped. To this end, we applied a single dPCA with the aim to decompose the variance attributed in CTOA-related, TA-related and RT-related components from the original dataset. In this case, trials were classified based on its CTOA (below the 40^th^ percentile or above the 60^th^ percentile of the CTOA distribution), its RT (below the 40^th^ percentile or above the 60^th^ of the RT distribution) and its TA (TA close (0°<TA≤6°), TA medium (6°<TA≤12°) and TA far (12°<TA<18°)), giving rise to 2×2×3 = 12 different potential classes where a trial can belong to.

Procedures to perform the dPCA analysis were performed using the MATLAB © (The Mathworkds Inc., Natick, Massachussetts) written scripts available from (14). Averaged MUA firing rate for each channel in each condition and each session were concatenated in a single dataset representing the entire recorded neural population, in the interval between -400 to 400 ms locked to the target onset. In each analysis, we used the decoding axis of each dPC assigned to each parameter as a linear classifier to decode trials belonging to the different categories (specific class for each parameter). To extract the statistical significance of this accuracy, we shuffled 100 times all available trials between classes and we thereby computed the distribution of classification accuracies expected by chance.

Since the coordinates of the components reflect the level of contribution to the activity of each neuron, the size of the dot product values between two components indicate that neurons that contribute to one component tend also to contribute to the other component. Therefore, the angle between two components can be interpreted as a marker of the functional overlapping between these components. When we applied the single dPCA analysis aimed to decompose the entire recorded population in latent variables associated to each parameter, we used the dot product between the first demixed principal components associated to each parameter to estimate the angle between these two components. For each pair of parameters, we measure the dot product between the subspaces obtained from first demixed principal component that maximally explained each parameter. Non-orthogonality is considered if the dot product between these two subspaces is greater than 3.3/N^1/2^, N being the number of total components considered in the decomposition (see (14) for details).

## Authors contributions

Conceptualization: J.A. and S.B.H.; Data Curation: J.L.A., F.D.B. and C.G.; Formal Analysis: J.L.A.; Funding Acquisition: S.B.H.; Investigation: J.L.A., F.D.B., S.B.H.H., E.A. and S.B.H.; Methodology: J.L.A., F.D.B., E.A. and S.B.H.; Resources: J.L.A. and S.B.H.; Supervision: S.B.H.; Validation: S.B.H; Visualisation: S.B.H; Writing-Original Draft: J.L.A. and S.B.H., Writing-review & editing: J.L.A., F.D.B., S.B.H.H., E.A., C.G. and S.B.H.

## Acknowledgments

J.A., C.G. were supported by ERC BRAIN3.0 # 681978 to S.B.H. and S.B.H.H. and F.D.B were supported by ANR-11-BSV4-0011 & ANR-14-ASTR-0011-01 to S.B.H. LABEX CORTEX funding (ANR-11-LABX-0042) from the Université de Lyon, within the program Investissements d’Avenir (ANR-11-IDEX-0007) operated by the French National Research Agency (ANR) to S.B.H. We acknowledge Inovarion for their fruitful scientific discussions. We also extend our thanks to Serge Pinède for hardware and software assistance and Jean-Luc Charieau and Fidji Francioly for animal care. F.B.is now affiliated to *Department of Physiology and Pharmacology, Sapienza University of Rome, 0018, Rome, Italy;* S.B.H.H. and C.G. are now affiliated to *Laboratory in Sensory Physiology, Otto Von Guericke University, 39120, Magdeburg, Germany;*

E.A. is now affiliated to *School of Innovation, Design, and Engineering, Mälardalen University, IDT, 721 23 Västerås, Sweden*.

## Data availability

The datasets supporting the current study have not been deposited in a public repository and are still under investigation. They are available from the corresponding author upon reasonable request.

## Code availability

Analysis were performed on Matlab, with custom codes, deposited onto a private github. They are available from the corresponding author upon reasonable request.

