## Supplementary Material for "The cost of multiplexing: PFC integrates multiple sources of information in non-orthogonal components accounting for behavioral variability"

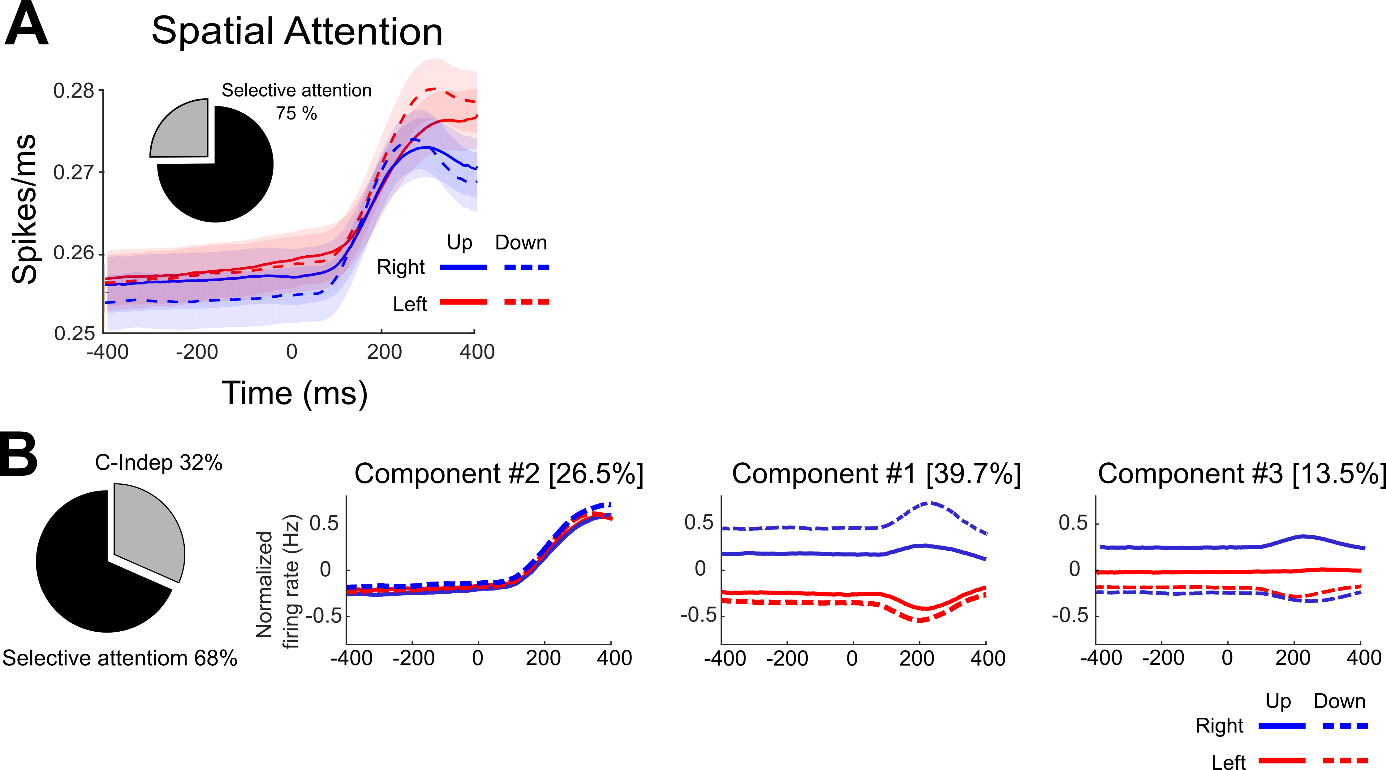

**Supplementary Figure 1. Spatial selectivity in the FEF.** (A) Mean MUA activity across all contacts (N = 848 electrodes) divided as a function of cued position (Continuous Blue: Right U; Dashed Blue: Right Down; Continuous Red: Left Up; Dashed Red: Right Down) locked to the target onset (-400 ms to 400 ms). Pie chart shows the proportion of position-cells (black, selective attention, 75%) in the recorded population. (B) Pie chart showing the proportion of variance explained by position (black, 68%) and position-independent (gray, 32%) in the recorded population. Projection of the firing rates averaged per each position onto the first three demixed principal components.

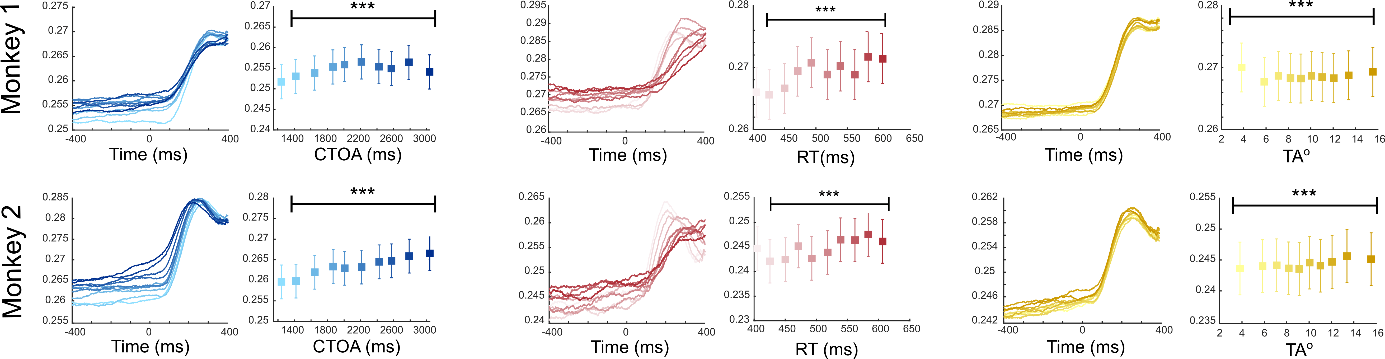

**Supplementary figure 2.** Time series for the MUA activity locked to the target onset for each monkey (M1, N = 480 contact; M2, N = 384 contacts). Tone colors represent the gradient of the parameter values (darker tones, higher values) for each of the parameters: CTOA (blue), RT (red) and TA (yellow). Boxplots showing the median and the interquartile range of the MUA activity before the target onset (-200 ms to 0) are also represented (Friendman test, *** p<0.001).

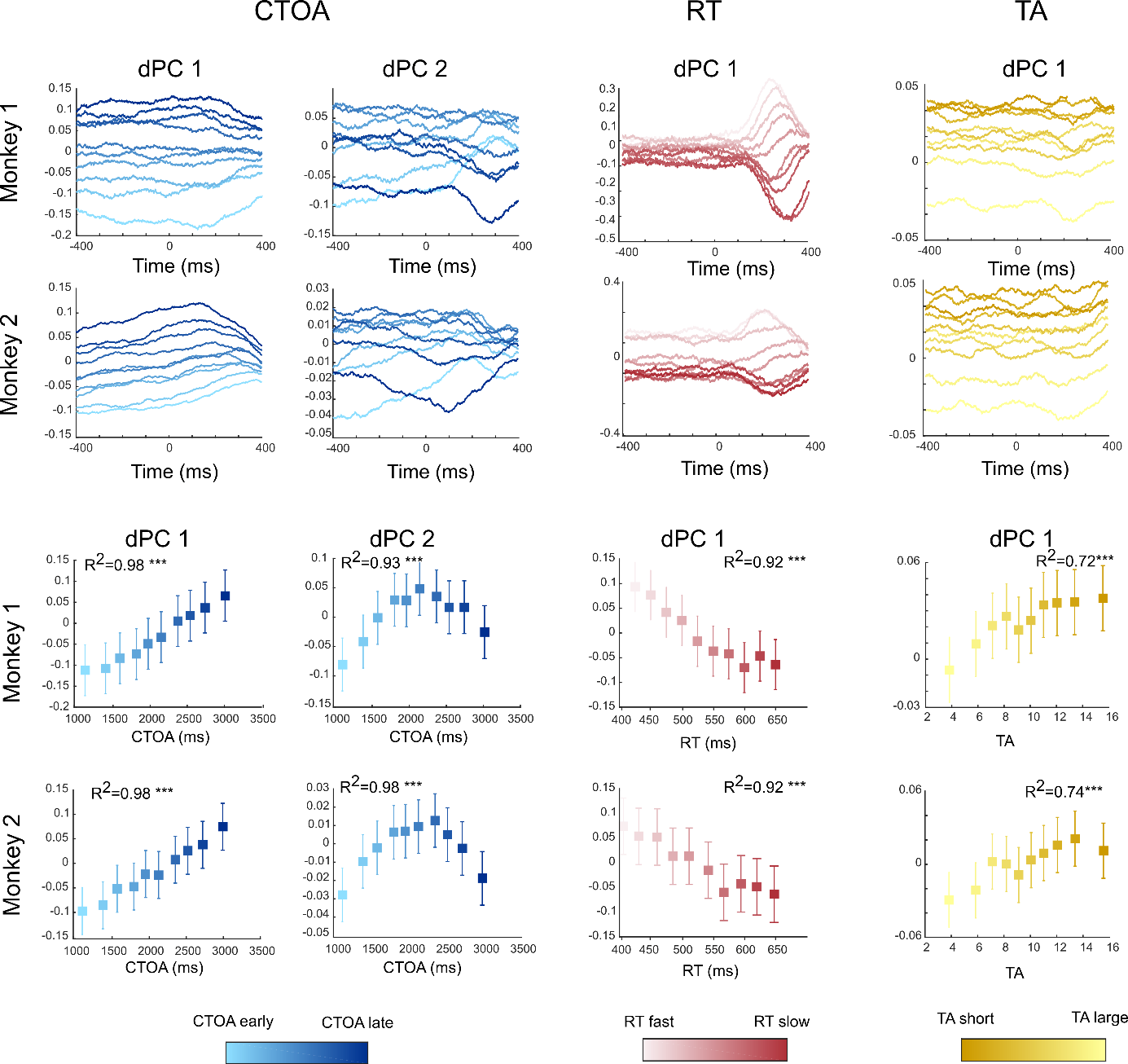

**Supplementary figure 3.** All as in figure 5.

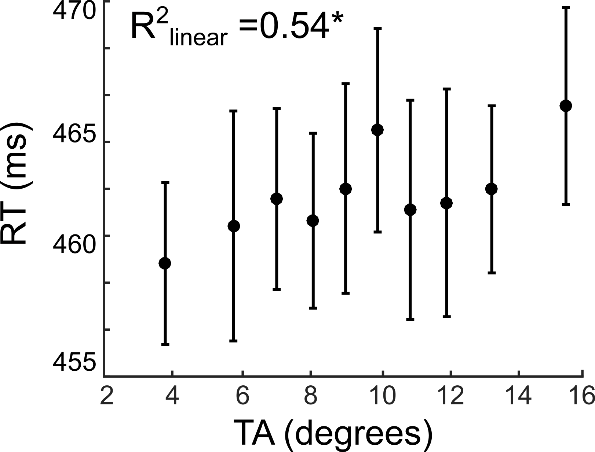

**Supplementary figure 4.** RT as a function of TA averaged across sessions. Each square represents, per each TA bin, the mean RT value. Bars represent the standard error of the RT measure across sessions. (* p < 0.05 linear regression)
